# Wing Hinge Dynamics Influence Stroke Amplitudes in Flapping Wing Insects: A Frequency Response Approach

**DOI:** 10.1101/2025.01.20.633950

**Authors:** Cailin Casey, Braden Cote, Chelsea Heveran, Mark Jankauski

## Abstract

Flapping wing insects leverage the dynamics of their compliant flight systems to reduce the energetic costs of flying. However, the extent to which the wing hinge dynamics contribute to the overall system dynamics remains unknown. Therefore, we developed an approach to (1) quantify the passive dynamic properties of the wing hinge, and (2) identify the resonant frequency of the isolated wing/wing hinge system. First, we measured the frequency response relating thorax deformation to wing stroke angle in sacrificed honeybees and army cutworm moths. Using these data, we developed a linear model of the flight system, which we then extended to incorporate nonlinear effects associated with large wing stroke angles. Our findings revealed that both species flap below the linear resonance of the wing hinge. At larger angles, nonlinear aerodynamic damping reduces the resonant frequency, causing both species to flap above wing hinge resonance. We discuss how wing-thorax coupling and muscle dynamics may cause the resonant frequency of the entire flight system to deviate from that of the wing/wing hinge system. Our estimates of wing hinge stiffness and damping provide quantitative parameters that can be incorporated into models of the insect flight system to enable more accurate predictions of resonance behavior.

## 1 Introduction

The flapping flight of small insects is widely recognized as one of the most energetically demanding forms of locomotion [1]. Unlike many birds, which glide to conserve energy [2], most insects must continuously beat their wings to counteract gravity, resulting in a high metabolic cost. Studies have shown that an insect’s metabolic rate during flight can reach levels 50-100 times higher than at rest [3]. To help offset this energetic expense, many insects have evolved a specialized flight mechanism known as indirect actuation, where the insect’s indirect flight muscles (IFMs) attach not to the wing base but to the thoracic exoskeleton [4, 5, 6]. When these muscles contract, they cause small deformations in the thorax, which are then amplified into large wing rotations through a complex linkage called the wing hinge [7, 8]. Small steering muscles within the wing hinge adjust its kinematics, allowing for quick, precise maneuvers during flight [9]. Indirect actuation likely helps reduce the energy required for flight by enabling the thorax to store and release energy at specific instances within the wing beat cycle, functioning as a spring-like structure [10].

Many insects are believed to flap at or near the resonant frequency of their flight system to reduce energetic cost [11, 12, 5]. This strategy is especially relevant for insects with asynchronous IFMs, which are believed to be activated at the resonant frequency of the system to which they are attached rather than the frequency of neurological activation [13]. Flapping at resonance reduces the force required to deform the thorax [14], which may lower muscle power output since power depends on both force and strain rate [3]. Deviating from this frequency could increase energetic demands, suggesting a trade-off between efficiency and maneuverability, particularly for insects that modulate wingbeat frequency during flight [12]. However, recent studies question the assumption that insects flap precisely at resonance. Models of hawkmoth flight indicate flapping occurs above the thorax’s resonant frequency, potentially improving control in response to aerodynamic disturbances [15]. Similarly, experiments on honeybees suggest their thorax resonance may exceed their wingbeat frequency, though artificial boundary conditions and low-amplitude excitations may inflate the measured resonance [16]. Other work suggests resonance in insect flight is not a single fixed state but may vary depending on whether displacement, velocity, or acceleration is used to define the system’s output [17]. However, even if insects do not flap exactly at resonance, they may still exploit the benefits of flapping near it, including energy recycling and vibration amplification.

Mathematical models are commonly used to estimate the resonant frequency of the flight system and to determine the energetic benefits of flapping near resonance. Qualitative models of the insect flight system combine effective stiffness into two spring-like elements: a ‘parallel’ stiffness element, representing the stiffness of the pleural wall in the thoracic exoskeleton as well as the passive stiffness of the IFMs and other thorax muscles, and a torsional ‘series’ stiffness element, representing the rotational stiffness of the pleural mechanism in the wing hinge [18, 17] (Fig. 1). However, most quantitative models idealize the wing hinge as a rigid lever mechanism [15], thereby neglecting the effective torsional stiffness and damping of the wing hinge. In reality, the wing hinge is a complex structure made up of stiff sclerites interspersed with softer connective membrane and muscle [7]. It is conceivable that the passive dynamics of the wing hinge contribute to the rotational kinematics of the flapping wing and consequently should be considered in modeling efforts of the flight system. Despite its potential importance, the dynamic properties of the wing hinge remain poorly characterized, and its mechanical role in flight is largely unquantified across insect species.

**Figure 1:**
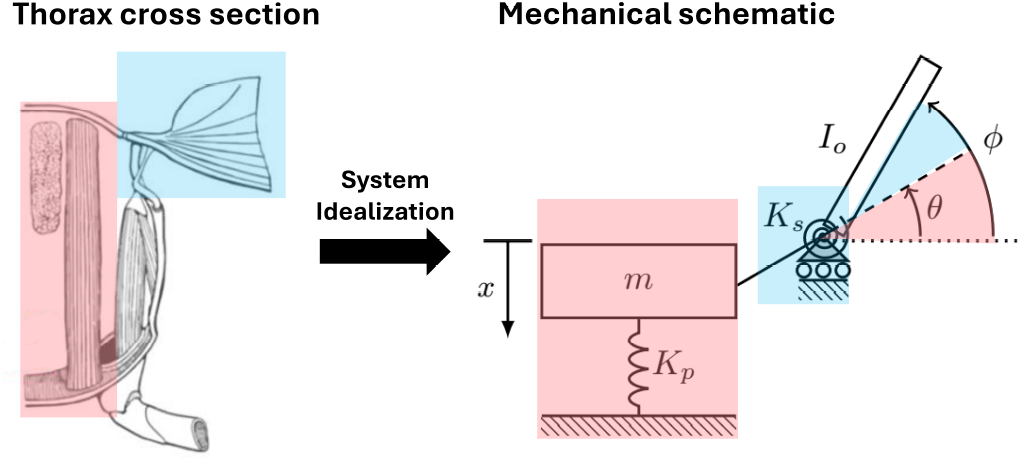
The thorax of insects utilizing indirect actuation (left) can be modeled as a two-degree-of-freedom system (right). In this model, parallel spring with stiffness *K*_*p*_ represents the combined effective stiffness of the thoracic exoskeleton and the passive stiffness of the IFMs, shown in red. A torsional spring with stiffness *K*_*s*_ represents the effective torsional stiffness of the pleural mechanism within the wing hinge, shown in blue. Parameters *m* and *I*_*o*_ denote the effective mass of the thorax and IFMs, and the mass moment of inertia of the wing about the wing hinge, respectively. Thoracic deformation, denoted as *x*, causes the wing to rotate by an angle *θ* under static conditions. However, under dynamic conditions, the stretching of the series spring modifies the wing’s rotation, resulting in an actual rotation angle *ϕ*. Left image adapted from [19], right image adapted from [18].

The objective of the present study is therefore to study the dynamics of the isolated wing hinge in order to (1) quantify its unknown dynamical properties, and (2) identify the resonant frequency of the isolated wing/wing hinge system and compare this to the insect’s flapping frequency. To this end, we developed a novel experimental approach to measure the frequency response relating thorax deformation to wing stroke angle in sacrificed insects. We considered two insect species: the army cutworm moth *Euxoa auxiliaris*, which is of the Lepidopteran order and has synchronous IFMs, and the honeybee *Apis Mellifera*, which is of the Hymenopteran order and has asynchronous IFMs. We restricted experimental thorax deformation and wing rotation amplitudes such that the flight system was excited primarily within the linear regime. We then utilized a simple mathematical model, populated in part by parameters estimated from the experiment, to estimate the nonlinear resonant frequency when the thorax and wings experience large amplitudes.

## 2 Methods

### 2.1 Specimen care and preparation

We selected the army cutworm moth (*E. auxiliaris*, n = 11) and the honeybee (honeybee, n = 10) as study species. We tested both forewings simultaneously for each subject, yielding a total of 22 wings tested for army cutworm moths and 20 wings tested for honeybees. Adult cutworm moths were collected from Powell, WY, and Bozeman, MT, and bred in captivity at Montana State University (Bozeman, MT, USA). Offspring from the F2 generation were used for the experiments. Moths were housed in a 1 m^3^ mesh cage at 20^*°*^C and fed daily with a 10% sugar solution delivered via soaked cotton balls. Specimens were tested within 10 days after emergence. Honeybees were captured near Montana State University and tested within 90 minutes of capture. All insects were euthanized in an ethyl acetate kill jar. Prior to testing, legs were removed using scissors. Thoracic scales on cutworm moths were gently removed with a damp paper towel.

### 2.2 Experimental setup and data collection

The experimental setup used to measure the frequency response between thorax deformation and wing stroke angle is shown in Fig. 2A. The thorax was situated between a fixed #2 set screw adhered to the dorsal thorax via cyanoacrylate and an electrodynamic shaker (2007E, The Modal Shop, Cincinnati, OH, USA), which compressed the thorax to induce wing rotation. For cutworm moths, the ventral thorax was adhered to a 3D-printed 15^*°*^ ramp with cyanoacrylate to orient the scutum normal to the set screw, consistent with similar experiments conducted on Lepidopteran species [20]. For honeybees, no ramp was needed, as the scutum was approximately normal to the set screw when the body was horizontal. To aid in tracking, a dot of nail polish was applied to the leading-edge vein (Fig. 2A). After setup, the insects remained in a compressed position for 10 minutes to allow the glue and nail polish to dry fully.

**Figure 2:**
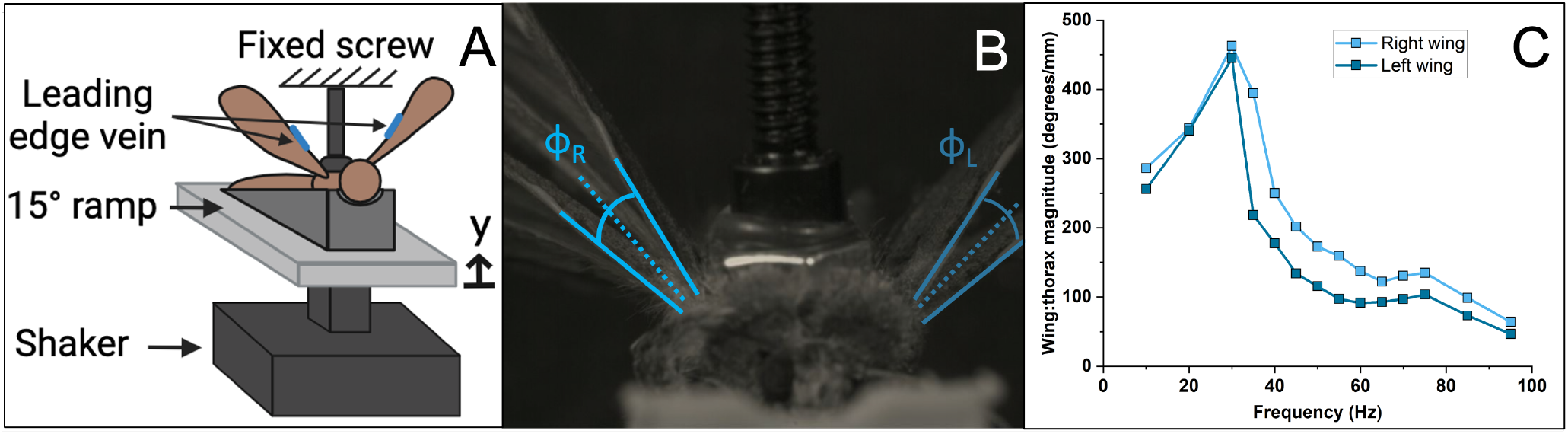
Experimental setup and data analysis. (A) Schematic of the thorax compression setup using a shaker, where *y* represents the shaker’s vertical displacement, assumed to coincide with the displacement of the thorax. (B) Overlaid frames captured by a high-speed camera showing the wing at its maximum upward and downward positions, where *ϕ* denotes the wing stroke angle amplitude (half the wing stroke). (C) Example frequency response function magnitudes showing the relationship between wing stroke angle and thorax displacement for the left and right wings.

The thorax was displaced harmonically via an electrodynamic shaker (2007E, The Modal Shop, Cincinnati, OH, USA). The input voltage to the shaker was modulated via a signal generator (SDG 1032X, Siglent, Solon, OH USA). A high-speed camera (Chronos 2.1, Kron Technologies, Burnaby, BC Canada) with macro lens (100mm f/2.8, Rokinon, New York, NY USA) was used to capture the wing rotation during testing. Frame rates of 2142 frames per second and 7135 frames per second were used for cutworm moth and honeybees, respectively. The camera was set to capture the projection of the wing stroke angle onto the insect’s transverse plane (Fig. 2B). We refer to this angle as the wing stroke angle hereafter. Note that in sacrificed insects, the plane from which wing stroke angle is referenced may differ from that observed in live insects because the steering muscles that modulate this plane are inactive. Insects were tested in a frequency range that encompassed their known flapping frequencies. Cutworm moth specimens were deformed at 14 discrete frequencies between 10 and 95 Hz (*in vivo* flapping frequency of 41 *±* 4.3 Hz [21]). Honeybee specimens were tested at 13 discrete frequencies between 50 and 350 Hz (*in vivo* flapping frequency of 230 Hz *±* 8.8 [22]).

Shaker displacement, assumed to coincide with thorax deformation at the location of the fixed screw, was also measured. For cutworm moths, an identical high speed camera with a 35mm lens (Rokinon, New York, NY USA) was synchronized with the first and positioned level with the shaker stage to measure vertical displacement. While similarly sized moths (*Helicoverpa Zea* and *Agrotis ipsilon*) had *in vivo* thorax displacements of 0.214 and 0.290 mm during tethered flight [23], we imposed smaller thorax compression amplitudes during this experiment (35 *±* 3.2*µ*m) to reduce the potential influence of nonlinearities on the frequency response, though we acknowledge that nonlinear aerodynamic damping may influence experimental results when the shaker excitation frequency approaches the resonant frequency of the wing/wing hinge system due to large wing stroke angular velocities. During testing, the shaker was started about a second before video recording commenced, so that only steady-state behavior was captured. Only the first wing cycle was analyzed for each frequency. The same procedure was followed for honeybees. A compression amplitude of 4.8*±* 0.35*µ*m was imposed on the thorax, which is lower than the thorax deformation amplitudes (*∼* 50*µ*m) observed during tethered flight of the similarly sized bumblebees [24]. For honeybees, such small shaker displacements could not be accurately observed using our high-speed videography equipment. We instead used a laser vibrometer (VibroGo VGO-200, Polytec, Baden-Württemberg, Germany) trained on the shaker stage supporting the insect to measure thorax displacement. Euthanasia to completion of testing took between 40 and 60 minutes for each insect. The exact measured shaker displacements for all subjects across all frequencies are shown in Fig. **??**.

### 2.3 Data analysis

Wing positions were tracked using DLTdv8 point software [25]. Four points were tracked for each specimen: one point at each root and one point each along the leading edge vein of the left and right wings. Wing stroke angle was calculated from the tracked points (Fig. 2C). The points on the leading edges were chosen in the first 50% of the wing length to minimize the potential confounding influence of wing deformation on rigid body wing stroke angle.

This study aimed to measure the wing stroke angle; however, wing pitching may have introduced some error into this measurement. The wing’s pitching rotation does not perfectly align with the leading-edge vein, as the pitching axis lies between the leading edge and the mid-chord line in hawkmoths [26] and perhaps in other insects as well. Consequently, pitch rotations could affect the calculated wing stroke angle. Despite this, we believe the error is small. High-speed video observations reveal relatively pitching at the small thorax displacements used in the experiment (though pitching tended to increase at the highest frequencies tested in honeybees; see supplementary material). Further, we tracked points along the leading edge vein near the wing root, where the pitching axis and leading edge vein are nearly coincident in some insects [27]. Multiple high-speed cameras would be required to capture the wing’s three-dimensional kinematics and fully decouple the wing’s stroke and pitch rotations, which was beyond the scope of the present work.

A Savitzky-Golay filter was applied to all time series data collected from the camera – both the left and right wing strokes, and the thorax displacement for cutworm moths. For cutworm moths, the filter had a frame size of 21 data points and an order of 3, meaning that every 21 points were fitted to a third degree polynomial. For honeybees, the frame size was smaller, 15 points, because there were fewer data points per curve at the highest frequencies. Representative wing kinematic data for one army cutworm and one honeybee (right wing angle vs. time for all frequencies, filtered and unfiltered) is shown in the supplementary material. The mean, median and interquartile range of wing kinematic data taken across all subjects at each excitation frequency is shown in supplementary material as well. The magnitude of the frequency response function was calculated by normalizing the wing stroke amplitude by the mean thorax compression amplitude at each frequency (Fig. 2C).

### 2.4 Frequency response curve fitting

Once the frequency response was determined experimentally under small amplitude assumptions, we fit a linear single-degree-of-freedom dynamical model relating thorax displacement *x* and stroke angle *ϕ*. The model is parameterized by natural frequency (*ω*_*n*_), damping ratio (*ζ*), and gain coefficient (*A*). The natural frequency and the damping ratio were estimated for each wing individually by curve fitting the magnitude data to a curve according to the following equation:

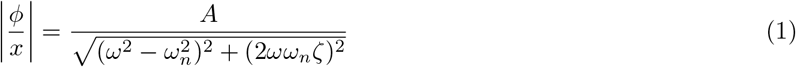

We used a nonlinear-least squares curve fitting approach (MATLAB version R2024b, function ‘lsqcurvefit’) to fit parameters. The lower and upper bounds for each parameter were specified a priori. The natural frequency *ω*_*n*_ was bounded between zero and approximately twice the species’ flapping frequency (0 to 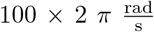for the army cutworm and 0 to 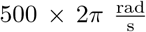 for the honeybee). For both species, the damping ratio *ζ* was bounded between 0 and 2, representing a range of values from undamped to overdamped. Because the gain coefficient *A* must be positive, the lower bound was set to 0 for both species. The upper bound for the gain coefficient was estimated by assuming a wing rotation amplitude of 1 rad and using *in-vivo* thorax deformation amplitudes approximated from the literature for similar species [23, 24] to approximate mechanical advantage *γ*. Then, we assumed the natural frequency to coincide with the reported insect flapping frequency. From this, we approximate a gain coefficient 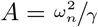. We then extended the upper bound of *A* beyond this value to allow for margin of error, resulting in an upper bound of 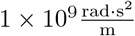for army cutworms and 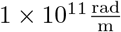for honeybees. The function tolerance was set to 1 *×* 10^*−*12^ and the optimality tolerance to 1 *×* 10^*−*12^. After curve fitting, we checked the results to ensure fitted parameters did not approach the specified parameter bounds. For both species, the frequency response magnitude data for the lowest and two highest experimental frequencies were excluded before performing curve fitting. This helped to reduce the influence of higher-order resonances on the accuracy of curve fitting around the wing hinge dominated resonance. We then discarded fits that had an *r*^2^ value less than 0.8, which reduced our effective sample size to *n* = 19 army cutworm wings and *n* = 17 honeybee wings (i.e., data from 3 wings discarded from each species). Poor fits generally occurred when there was no identifiable peak within the frequency range tested.

### 2.5 Modeling

Aerodynamic damping acting on the wing at large stroke amplitudes introduces a quadratic nonlinearity. As a result, we expect the shape and resonance of the frequency response to be affected by the amplitude of the input thorax deformation [15]. We developed a mathematical model to estimate how the frequency response relating thorax displacement to wing stroke angle changes at large input thorax deformation amplitudes (Fig. 3).

**Figure 3:**
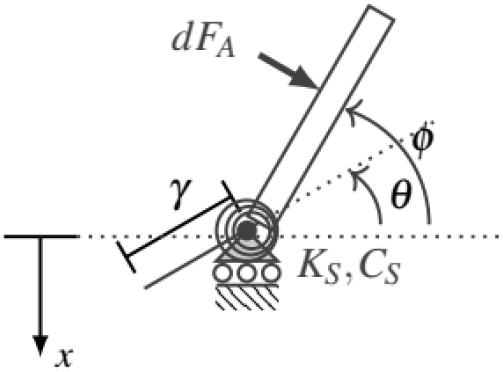
Simple mechanical model of the insect flight system. *x* denotes thorax deformation, which algebraically relates to the angle *θ* prescribed at the base of the wing by *θ* = ^*x*^, where *γ* is the transmission ratio. The actual wing angle is denoted *ϕ*. The effective torsional stiffness and damping at the wing hinge are *K*_*s*_ and *C*_*s*_, respectively. *dF*_*a*_ is a differential aerodynamic force acting on the wing, which is integrated over the wing surface to yield the total aerodynamic force and moment.

Suppose that the thorax deformation is *x*, which is prescribed within the experiment and consequently can be neglected as an independent degree-of-freedom. The thorax connects to the base of the wing via a rigid lever. Thorax deformation *x* is algebraically related to the angle prescribed at the base of the wing *θ* by 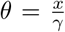, where *γ* is the wing-thorax transmission ratio. Due to local compliance, the wing is permitted to rotate beyond what is prescribed at the wing base by the thorax deformation. Actual wing rotation is defined as *ϕ*. The aerodynamic moment acting on the wing is determined through a blade element approach [28, 29]. We use a moment balance about the wing base to derive the equation of motion governing *ϕ* as

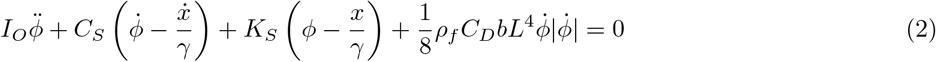

where *K*_*s*_ and *C*_*s*_ are the torsional stiffness and damping at the wing base, respectively, *I*_*o*_ is the wing’s mass moment of inertia about its base, *ρ*_*f*_ is fluid density, *b* is the wing’s chord width (assumed constant), *L* is the wing length and *C*_*D*_ is a drag coefficient.

The model parameters were populated using a combination of anatomical measurements, literature, and experimental data (Tab. 2). We modeled the wing as a thin, uniform, rigid rectangle. The wing’s mass moment of inertia is 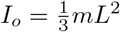, where *m* is the wing’s mass. The forewings were massed (AS 60/220.R2 Plus, Radwag, Radom Poland) and their length and maximum width were measured using ImageJ [30]. All wing measurements were averaged from the left and right wings of two individuals and are for the forewings only. Air density was based on its density at standard temperature and pressure. We considered a drag coefficient of 2, which lies between the minimum (0.4) and maximum (3.4) values reported in [31]. While the drag coefficient varies temporally with angle of attack in real flapping wings, our models assume single degree-of-freedom rotation and thus a constant angle of attack. The drag coefficient is therefore treated as constant. Although a value closer to the upper bound might better reflect the primarily normal induced flow expected under the simplified model kinematics, we selected a value of 2 to avoid overestimating the influence of drag relative to that observed in real insect flight. The remaining model parameters were populated using fitted frequency response data. The torsional stiffness and damping at the wing base are calculated based on the system’s fitted natural frequency and damping ratio (Eq. 1) and the wing’s idealized moment of inertia. Despite the small amplitude excitation considered in the experiment, nonlinear aerodynamic may increase the perceived damping in the experimental data. Thus, the torsional damping calculated from the fitted experimental data likely represents an overestimate of the torsional damping in the underlying linear system. The transmission ratio is inferred from the static gain of the frequency response.

Thorax displacement, including displacement amplitude and frequency, served as the model input. Thorax displacement was assumed harmonic and its amplitude was varied in both insects such that the resulting wing stroke amplitudes spanned from the experimental range to the *in vivo* range. This yielded displacement amplitudes of 35, 100, 250 and 500 *µ*m for cutworm moth and 5, 25, 75, 150 *µ*m for honeybees. The model was used to find the steady-state wing stroke amplitude over a frequency range similar to the experimental range (1 to 100 Hz for cutworm moth and 1 to 350 Hz for honeybees). For each frequency, the model was solved at 1000 time points over 20 wingbeats to reach steady state using ODE45 (MATLAB version R2024b). The stroke amplitude of the final wingbeat was used to calculate the ratio of wing stroke amplitude to thorax displacement amplitude.

## 3 Results

### 3.1 Experimental results

The frequency response magnitudes for each insect, separated by left and right wings, are shown in Fig. 4A,B,D,E. Individual magnitude curves were fitted via Eq. 1 to identify the gain coefficient, natural frequency, and damping ratio for each insect. Each quantity was subsequently averaged to obtain the data shown in Tab. 1. We then used the average data from Tab. 1 to calculate a frequency response magnitude representative of all subjects within a species, shown in Fig. 4C,F. According to the representative frequency response magnitude plots, the resonant frequency is about 52.7 Hz in army cutworms and 219.7 Hz in honeybees. This is lower than the system natural frequencies reported in Tab. 1, which is expected in systems with modest damping. Both insects displayed relatively low damping ratios. As noted previously, the damping ratios reported here likely overestimate the real values, since nonlinear aerodynamic damping may influence experimental measurements even at relatively low wing stroke amplitudes. Representative videos of the wing stroke angle at pre, near and post resonance conditions for both species are shown in supplementary material.

**Figure 4:**
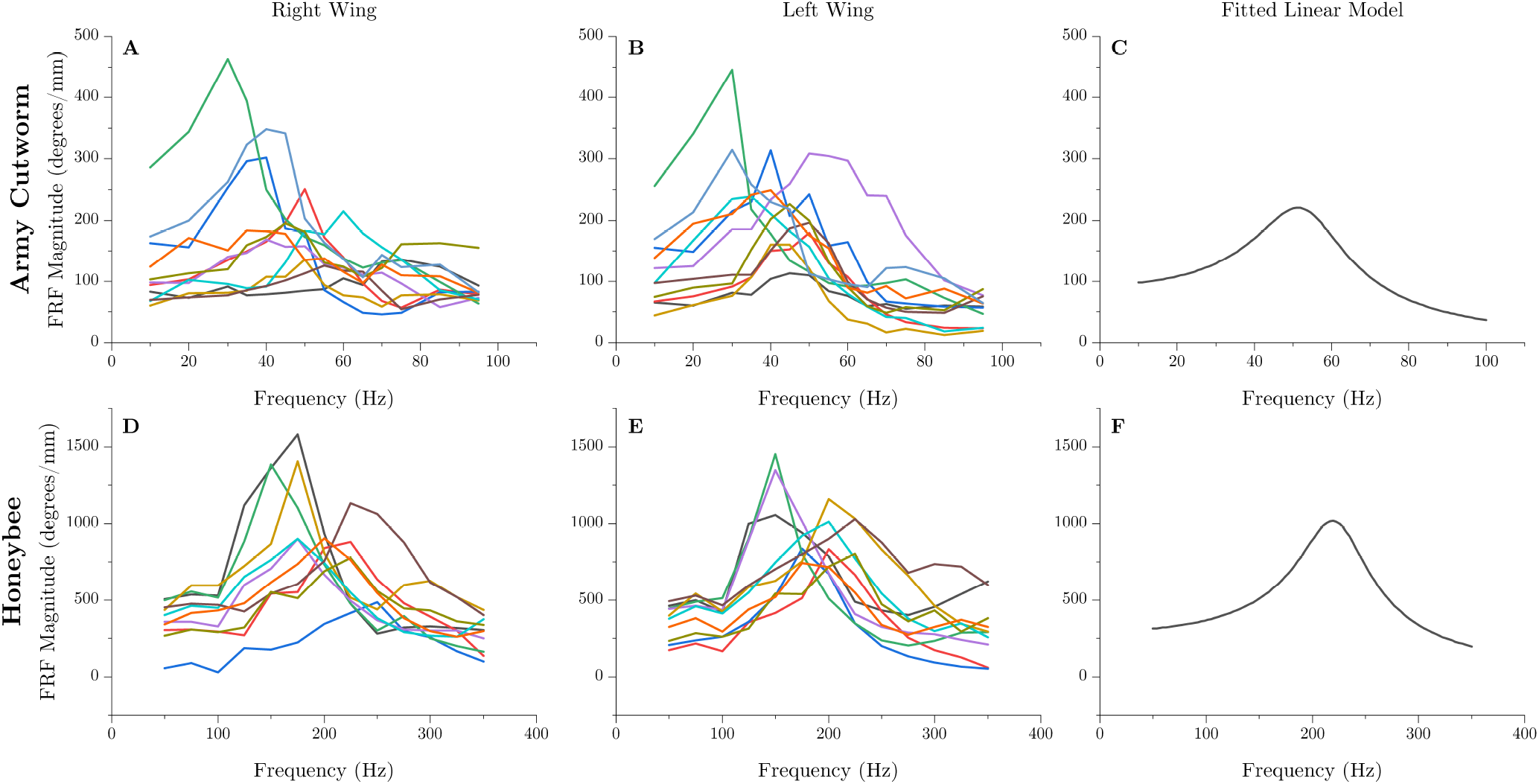
Frequency response magnitude as a function of excitation frequency for all subjects. The right-hand panels plot Eq. 1 populated with the average natural frequencies, damping ratios and gain factors reported in Tab. 1, which represents a typical frequency response magnitude for each species.

**Table 1:** Linear frequency response parameters. Wing responses were averaged across all individuals (mean*±* standard deviation). Left and right wings are treated as individual trials.

| Description | Variable | Army Cutworm | Honeybee | Units |
| --- | --- | --- | --- | --- |
| Sample size | $n$ | 19 | 17 | - |
| Gain Coefficient | $A$ | $(1.91 \pm 0.66) \times 10^2$ | $(1.03 \pm 0.42) \times 10^4$ | $\frac{\text{rad} \cdot \text{s}^2}{\mu\text{m}}$ |
| Natural Frequency | $\omega_n$ | $53.98 \pm 9.78$ | $224.14 \pm 30.33$ | Hz |
| Damping Ratio | $\zeta$ | $0.22 \pm 0.06$ | $0.15 \pm 0.04$ | - |

### 3.2 Modeling results

The parameters used in the numerical models are listed in Tab. 2. Figure 5 illustrates the frequency response amplitude and response stroke angle amplitude for varying levels of thorax compression in both species. For the army cutworm, at low thorax deformation amplitudes, the flapping frequency is lower than the flight system’s resonant frequency. However, as the thorax deformation amplitude increases, the flapping frequency surpasses the resonant frequency. In the case of the honeybee, the flapping frequency matches the flight system’s resonant frequency at small thorax deformation amplitudes. With greater thorax deformation, the flapping frequency exceeds the resonant frequency. In both species, the observed reduction in resonant frequency at high thorax deformation amplitudes is caused by nonlinear aerodynamic damping. Real flapping wing insects exhibit stroke amplitudes of approximately 45 - 60^*°*^ [22, 32], which the model predicts occurs only near the highest amplitudes of thorax deformation considered.

**Figure 5:**
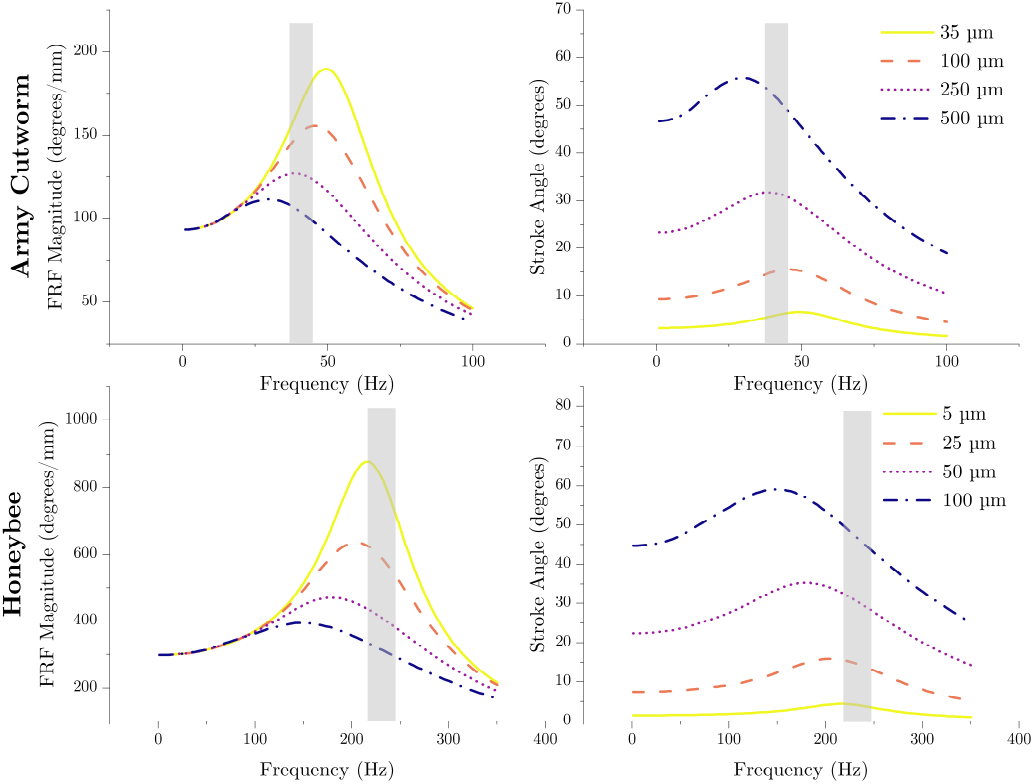
Nonlinear modeling predictions of the frequency response magnitude and wing stroke angle as a function of flapping frequency for both species and various thorax deformation amplitudes. The gray regions indicate the reported flapping frequency range for both species.

**Table 2:** Model parameters.

| Symbol | Equation | Army Cutworm | Honeybee | Units |
| --- | --- | --- | --- | --- |
| $\zeta$ | | 0.26 | 0.17 | - |
| $\omega_n$ | | 50 | 204 | Hz |
| $L$ | | 19 | 8.1 | mm |
| $b$ | | 7.0 | 2.3 | mm |
| $m$ | | 1.9 | 0.15 | mg |
| $\rho_f$ | | 1.25 | 1.25 | $\frac{\text{kg}}{\text{m}^3}$ |
| $C_D$ | | 2 | 2 | - |
| $I_O$ | $\frac{1}{3}mL^2$ | $2.28 \times 10^{-10}$ | $3.28 \times 10^{-12}$ | $\text{kg} \cdot \text{m}^2$ |
| $K_S$ | $\omega_n^2 I_O$ | $2.63 \times 10^{-5}$ | $6.51 \times 10^{-6}$ | $\frac{\text{N} \cdot \text{m}}{\text{rad}}$ |
| $C_S$ | $2\zeta\omega_n I_O$ | $3.41 \times 10^{-8}$ | $1.39 \times 10^{-9}$ | $\frac{\text{N} \cdot \text{m} \cdot \text{s}}{\text{rad}}$ |
| $\gamma$ | $\frac{\omega_n^2}{A}$ | $6.14 \times 10^{-4}$ | $1.92 \times 10^{-4}$ | $\frac{\text{m}}{\text{rad}}$ |

## 4 Discussion

Flapping wing insects exploit the dynamics of their flexible flight system to reduce the energetic costs of flight. Recent models of the flight system lump this flexibility into a parallel spring, representative of the exoskeleton and passive stiffness of the IFMs and other thorax muscles, and a series spring, representative of the torsional stiffness of the wing hinge [18]. While experimental studies have addressed the mechanics of the thorax, the dynamics of the wing hinge has received comparatively less attention. The objective of this study was to determine the mechanical properties and resonant frequency of the isolated wing/wing hinge system in army cutworm moths and honeybees using a frequency response-based approach.

The resonant frequency of the wing/wing hinge system was in proximity to the flapping frequency for both the honeybee and army cutworm. Specifically, the army cutworm flaps at approximately 41 Hz [21] and the honeybee at about 230 Hz [22], while small amplitude experiments identified linear resonant frequencies of roughly 53 Hz and 220 Hz respectively (Fig. 4). This implies that both species flap below the linear resonant frequency of the wing/wing hinge system. As wing stroke amplitudes increase to *in vivo* values, mathematical models predict that both species flap at frequencies above nonlinear resonance of the wing/wing hinge system (Fig. 5). However, the extent to which the resonant frequency of the isolated wing/wing hinge system reflects that of the full flight system remains uncertain. Interpreting these results in the broader context of insect flight requires consideration of additional factors, including interactions with the thorax and inclusion of muscle dynamics, which are detailed here.

### 4.1 Wing-Thorax Coupling

First, coupling between the wing/wing hinge system and the flexible thorax, as well as thorax mediated wing-wing coupling, must be considered when evaluating the resonant frequency of the entire flight system. Experimentally, we prescribed thorax deformation rather than the forces acting on the thorax. This restricts backward coupling between the wing and the thorax (that is, the thorax motion influences the wing, but wing dynamics cannot influence the deformation of the thorax at the parallel spring location) as well as wing-wing coupling through the thorax. The wings can therefore behave as independent oscillators, which is why the left and right wings of an individual insect may have different resonant frequencies (Fig. 4). Experimentally, the left-right resonant frequencies may differ if the set screw actuating the thorax is slightly misaligned, or if there is difference in initial wing orientation when the insect is sacrificed. Previous studies in flies support the idea that the wings can behave as independent oscillators, where disruption of thorax-mediated wing-wing coupling led to disparate flapping frequencies between wing pairs [11]. Should thorax flexibility be considered, we expect that the effective resonant frequency of the wing/wing hinge system will be reduced due to a compliant boundary condition.

The degree to which thorax coupling will influence the flight system resonant frequency depends on the proximity of the resonant frequencies of the isolated wing/wing hinge and the isolated thorax. If the resonant frequency of the isolated thorax is considerably higher, the coupling will be minimal, and the resonant frequency of the wing/wing hinge would closely reflect that of the entire flight system. Experimental studies on the *M. sexta* suggest that the thorax does not have a resonant frequency in a range of up to about 3 times the insect’s flapping frequency [10], though it is unclear if this will generalize to other species.

Mechanically, the resonant frequency of the isolated thorax depends on the stiffness of the thoracic exoskeleton and the passive stiffness of the IFMs and other thoracic muscles as well as the effective mass of these components. The effective mass of the thorax and IFMs has been shown to be small compared to the rotational inertia of the wings in many species [33]; thus, the thorax behaves primarily as a spring-damper. Studies have characterized the effective passive stiffness of the the thorax and IFMs, which can be thought of as *K*_*p*_ (Fig. 1). It can be shown that the stiffness ratio 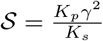, where *K*_p_ is the parallel stiffness, *K*_s_ is the series stiffness, and *γ* is the transmission ratio. The latter two quantities are reported in Tab. 2. We did not calculate *K*_*p*_ in the present study for the honeybee or army cutworm moth. We instead estimated *K*_*p*_ for the honeybee using values for *Bombus centralis* (*K*_*P*_ = 973 N/m) and for the army cutworm using values for *Agrotis ipsilon* (*K*_*P*_ = 481 N/m) because they are of similar size and have the same muscle type as the test species [23]. Using these values, we arrive at stiffness ratios of *S* = 6.9 for the army cutworm and *S* = 5.5 for the honeybee. Thus, the relative stiffness of the series spring appears to be similar in the honeybee compared to the army cutworm – and for both species, the relative stiffness of the parallel spring is higher than that of the series spring. If we assume that the effective mass of the IFMs and thoracic exoskeleton is small compared to the rotational inertia of the wing, which is consistent with the findings in [33], a stiffness ratio greater than one implies that the resonant frequency of the isolated thorax is higher than that of the wing/wing hinge system.

### 4.2 Muscle Dynamics

Muscle dynamics may also influence the system’s effective resonant frequency, particularly in the case of the honeybee with asynchronous IFMs. Asynchronous muscle models suggest that the force they produce is proportional to the muscle strain rate processed through a second-order low-pass filter [34]. Because muscle forces are a function of muscle strain rate and not simply time, they must be considered among the flight system’s dynamical state and consequently influence the system’s resonant frequencies. All experiments in this work considered sacrificed insects with inactive IFMs; the present experiment captures only the *mechanical* resonance of the wing hinge. Inclusion of muscle dynamics may lead to a *musculo-mechanical* resonance, which may occur at a frequency different than the mechanical resonance depending on the specific muscle properties. Although we expect the muscle dynamics influence the resonant frequency more in insects with asynchronous IFMs, it is possible that muscle dynamics influence resonant frequencies in insects with synchronous IFMs as well. Recent evidence in the hawkmoth *M. sexta* show that their synchronous flight muscles exhibit delayed stretch activation, a property usually attributed to asynchronous muscles [35]. Consequently, the possibility of musculo-mechanical resonance should not be ignored in insects with synchronous IFMs.

### 4.3 Resonance Types

Multiple types of resonance may exist within the flight system [17]. In our experiment, we measured the frequency response relating thorax deformation to wing stroke angle, which are both position variables. However, the resonant frequency will be different depending on if angular position, velocity or acceleration is considered as the kinematic output. A displacement-displacement frequency response typically has a resonant frequency lower than the system’s natural frequency when damping is present [36]. By contrast, a displacement-velocity frequency response has a resonant frequency nearly equivalent to the system’s natural frequency and a displacement-acceleration higher than the system’s natural frequency [36]. To explore these alternative resonant states, we plot the response angle, angular velocity and angular acceleration determined via the nonlinear model for various thorax deformation amplitudes in Fig. 6. At the largest thorax deformation amplitudes considered, the flapping frequency of the army cutworm falls between the angular displacement and angular velocity resonant frequencies. For the honeybee, the flapping frequency is nearly coincident with the angular velocity resonance. It is possible that differing species, particularly those with different types of IFMs, leverage different types of resonance to realize flight.

**Figure 6:**
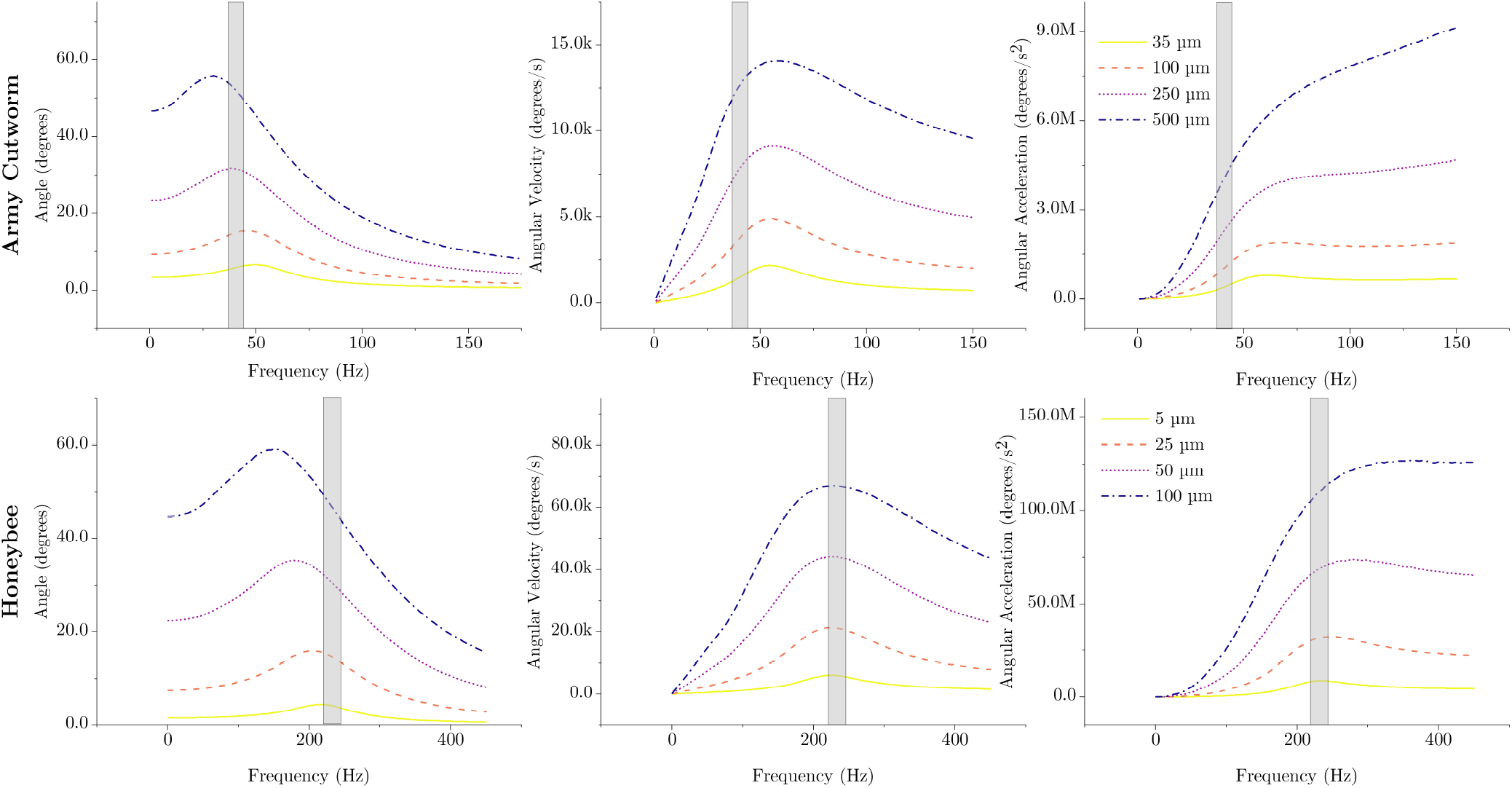
Mathematical model predictions of angular displacement, velocity and acceleration responses for various thorax deformation amplitudes. The gray box indicates the approximate flapping range for both species.

### 4.4 Future Outlook

While this study provides new insights into the mechanical properties and resonant behavior of the isolated wing/wing hinge system, several questions remain. Future work should investigate the dynamics of the complete flight system, incorporating both thorax-wing coupling and the contributions of active muscle forces. In particular, experiments involving anesthetized insects could help quantify how muscle activation, especially the delayed stretch activation observed in both asynchronous and some synchronous IFMs [35], influences the system’s effective resonance. Further, the present study relied on simplified planar kinematics and small amplitude excitation. Threedimensional measurements of wing motion, including pitch and stroke coupling, would provide a more complete understanding of wing-hinge dynamics during natural flight. This may be particularly important for capturing nonlinear effects that emerge at larger stroke amplitudes. From a modeling perspective, integrating these factors into a multi-degree-of-freedom model that includes thoracic compliance, bilateral wing coupling, and active muscle dynamics could improve our ability to predict flight performance across species. Finally, expanding this framework to a broader range of insect taxa would reveal whether tuning wingbeat frequency near a specific resonance is a universal strategy or one that varies depending on evolutionary or ecological constraints.

## Author Contributions

Conceptualization - MJ; Data curation - CC, MJ; Formal Analysis - BC, CC, MJ; Funding Acquisition - CC, MJ; Investigation - CC; Methodology - CC, CH, MJ; Project Administration - CH, MJ; Software - CC, MJ; Supervision - CH, MJ; Validation - CC; Visualization - CC; Writing, original draft - CC, MJ; Writing, review & editing - BC, CC, CH, MJ

## Acknowledgments

We would like to thank Taylor Kennedy for raising the *E. auxiliaris* specimens. We would also like to thank the anonymous reviewers for their constructive feedback.

## Data Accessibility

The data and models supporting this work are available at https://zenodo.org/records/14675752.

